# A unified model for the biogenesis of mitochondrial outer membrane multi-span proteins

**DOI:** 10.1101/2021.12.14.472550

**Authors:** Jialin Zhou, Martin Jung, Kai S. Dimmer, Doron Rapaport

## Abstract

The mitochondrial outer membrane (MOM) harbors proteins that traverse the membrane via several helical segments, so-called multi-span proteins. Two contradicting mechanisms were suggested to describe their integration into the MOM. The first proposes that the <u>m</u>itochondrial <u>im</u>port (MIM) complex facilitates this process and functions as an insertase, whereas the second suggests that such proteins can integrate into the lipid phase without the assistance of import factors in a process that is enhanced by phosphatidic acid. To resolve this discrepancy and obtain new insights on the biogenesis of these proteins, we addressed this issue using yeast mitochondria and the multi-span protein Om14. Testing different truncation variants, we show that only the full-length protein contains all the required information that assure targeting specificity. Employing a specific insertion assay and several single and double deletion strains, we show that neither the import receptor Tom70 nor any other protein with a cytosolically exposed domain have a crucial contribution to the biogenesis process. We further demonstrate that Mim1 and Porin are required for optimal membrane integration of Om14 but none of them is absolutely required. Unfolding of the newly synthesized protein, its optimal hydrophobicity, as well as higher fluidity of the membrane dramatically enhanced the import capacity of Om14. Collectively, our findings suggest that MOM multi-span proteins can follow different biogenesis pathways in which proteinaceous elements and membrane behavior contribute to a variable extent to the combined efficiency.

**Summary:** Zhou et al. provide new insights to the biogenesis of mitochondrial outer membrane proteins. They demonstrate that such proteins can follow various routes where both proteinaceous elements and membrane behavior regulate the efficiency and specificity of this process.

## Introduction

Mitochondria are double-membrane organelles, which according to the endosymbiotic hypothesis, descended from engulfed aerobic prokaryotes. As the main power house of eukaryotic cells, mitochondria possess about 800 proteins in yeast and 1500 in humans, of which 99% are encoded in the nucleus, synthesized by cytosolic ribosomes, and then directed to their specific mitochondrial sub-compartment (Neupert and Herrmann, 2007; Walther and Rapaport, 2009; Drwesh and Rapaport, 2020; Pfanner et al., 2019).

While the mitochondrial inner membrane sets a border between the intermembrane space (IMS) and the matrix, the mitochondrial outer membrane (MOM) constitutes a barrier to separate the organelle from the rest of the cellular environment. Transmembrane proteins residing in the MOM play a critical role in many mitochondrial activities such as protein biogenesis, the formation of contact sites with other organelles, maintenance of mitochondrial morphology, as well as modulation of mitochondrial motility (Friedman et al. 2011; Hermann et al. 1998; Cohen et al. 2009; Sogo and Yaffe 1994). Dysfunction of these proteins may lead to failure in the aforementioned processes and even to severe human diseases such as neurodevelopmental disorders (Ghosh et al. 2019; Wei et al. 2020). Considering their importance, it is critical to understand the biogenesis of MOM proteins, including their targeting to the organelle, membrane integration, and final maturation into a stable and functional form. In the past decades, some insights on targeting and integration of multi-span helical MOM proteins have been obtained (Becker et al., 2011; Sauerwald et al., 2015; Vögtle et al., 2015; Papić et al., 2011; Coonrod et al., 2007; Doan et al., 2020; Sinzel et al., 2017). The mitochondrial outer membrane receptor Tom70 has been found to support the targeting process by either directly recognizing the precursors of multi-span proteins or indirectly by serving as a docking site for chaperones that bind to and stabilize such newly synthesized proteins (Backes et al., 2021; Doan et al., 2020; Otera et al., 2007; Papić et al., 2011; Becker et al., 2011). In yeast, two additional MOM proteins, Mim1 and Mim2, are also involved in the biogenesis of multi-span proteins before and during their integration into the MOM (Becker et al., 2011; Papić et al., 2011; Dimmer et al., 2012). However, the exact role of Mim1/2 is yet to be elucidated as it was reported that newly synthesized Ugo1, a multispan MOM protein, can be inserted *in vitro* into pure lipid vesicles in a process that was enhanced by elevated levels of phosphatidic acid within the lipid bilayers (Vögtle et al., 2015). Similarly unclear are the initial cytosolic steps in the biogenesis of such multi-span proteins as well as the targeting signals that assure their specific sorting to the organelle.

To shed light on these understudied processes, we decided to use the multi-span MOM protein Om14 as a model protein. Om14 is one of the most abundant MOM proteins and it contains three predicted α-helical transmembrane domains (TMDs), exposing the N-terminus towards the cytosol and the C-terminus towards the IMS (Burri et al., 2006). The function of Om14 is not resolved yet. On the one hand it was reported to form a complex with the MOM proteins VDAC (Porin in yeast) and Om45 and be involved in the import of metabolites (Lauffer et al., 2012), whereas on the other hand Om14 was suggested to function as a receptor for cytosolic ribosomes (Lesnik et al., 2014). Although it is still unclear whether the targeting of Om14 is promoted by the Tom70 receptor, an interaction of newly synthesized Om14 molecules and cytosolic chaperones like Hsp70, Hsp90 and Hsp40 cochaperones such as Ydj1 and Sis1 was reported (Jores et al., 2018). Similar to Ugo1, the biogenesis of Om14 has been suggested to be dependent on Mim1 (Becker et al., 2011). Additionally, the cardiolipin level in the mitochondria affects the biogenesis of Om14 (Sauerwald et al., 2015). Even though some common factors like Tom70 and the MIM complex seem to support the biogenesis of both Om14 and Ugo1, it is currently unclear what are the contributions of other features like folding of the protein, its hydrophobicity, and the behavior of the lipid phase. It might well be that, similarly to the variety of pathways described for single-span proteins (Vitali et al., 2020), multi-span proteins follow rather individual routes.

To address these issues, we investigate here cis and trans elements that might impact on Om14 biogenesis. Our findings indicate that the targeting and membrane integration of Om14 involve cis elements like hydrophobicity of the putative TMDs and an unfolded state as well as multiple trans factors like Tom70, Mim1, and Porin, together with the fluidity of the membrane. The comprehensive evaluation of the modulatory effects of all these factors provides a new concept for our understanding of the biogenesis of MOM proteins.

## Results

### Multiple targeting signals within Om14 collectively contribute to its mitochondrial location

As the initial step of our efforts to dissect the biogenesis pathway of Om14, we asked which parts of the molecule serve as a mitochondrial targeting signal. To address this question, we fused GFP to full-length OM14 or to three segments including each one of the three putative TMDs of Om14 and its flanking regions (Fig. 1A). To minimize interference from the GFP module, the fusion proteins were constructed according to a suggested topology model such that the GFP moiety does not have to cross the membrane (Burri et al., 2006). Next, we investigated whether one of the individual TMDs or a truncated protein containing two of them are sufficient for mitochondrial targeting. To this end, we examined the localization of these truncated variants by fluorescence microscopy (Fig. 1B). As expected, the full-length Om14 showed a clear co-localization with mitochondrial targeted RFP (Mito-RFP). While all the truncated constructs display clear mitochondrial signal, all three constructs with a single TMD displayed also some GFP-stain in the cytosol (Fig. 1B). This cytosolic signal can result, at least partially, from proteolytic cleavage of the OM14 portion and a cytosolic location of the remaining GFP moiety. We also observed a possible ER localization of the construct GFP- Om14(1-62) as indicated by the peri-nuclear fluorescent signal (Fig. 1B, marked with an arrow). To further study the intracellular location of the GFP-tagged fragments, we performed a sub- cellular fractionation procedure and analyzed the localization of the different constructs via immunodecoration with anti-GFP antibody (Fig. 1C). In line with our fluorescence microscopic findings, the full-length Om14 was detected solely in the mitochondrial fraction, while all the truncated constructs were detected primarily in the mitochondrial fraction with a variable subpopulation mis-localized to the ER. In addition, a band slightly smaller than the intact GFP- Om14(1-62) construct, which is caused probably by a cleavage of several amino acids at the C-terminus of this variant, was observed in the whole cell lysate and cytosol fractions but not in the mitochondrial one (Fig. 1C). This indicates that amino acid residues at the C-terminus of this construct are important for mitochondrial targeting. In two cases (GFP-Om14(1-62) and GFP-Om14(96-134)), a clear band in the cytosolic fraction at the expected size of GFP alone could be detected (Fig. 1C, “*”). This observation supports our proposal that a major portion of the cytosolic GFP fluorescence signal might be related to degradation products.

**Figure 1.**
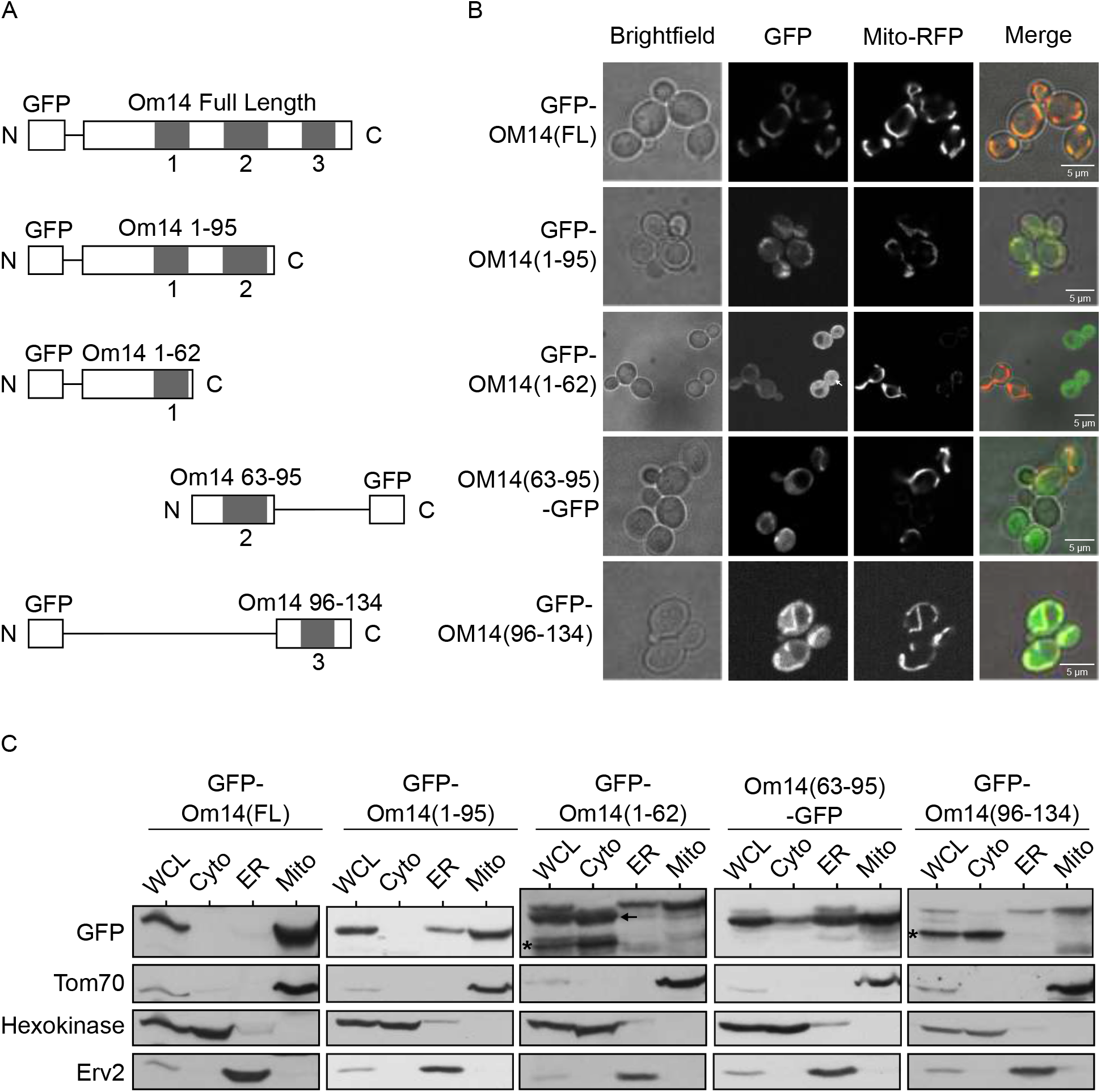
Om14 harbors multiple mitochondrial targeting signals. **(A)** Schematic representation of GFP fusion proteins of full length Om14 and its truncated variants. **(B)** Localization of Om14 constructs as visualized by fluorescence microscopy. RFP targeted to mitochondria (Mito-RFP) was used as a marker for mitochondrial structures. **(C)** Cells expressing the various Om14 variants were subjected to sub-cellular fractionation. Fractions corresponding to whole cell lysate (WCL), cytosol (Cyto), microsomes (ER), and mitochondria (Mito) were analyzed by SDS-PAGE and immunodecoration with the indicated antibodies. Tom70, Hexokinase, and Erv2 were used as markers for mitochondria, cytosol, and microsomes, respectively. Asterisk marks the GFP moiety alone whereas an arrow indicates a slightly processed form of GFP- Om14(1-62).

The capacity of the single TMDs to associate with mitochondria was supported by *in vitro* import assays where radiolabeled constructs containing the single TMD were imported into isolated mitochondria (Fig. S1). Interestingly, this import was hardly dependent on the presence of Mim1, pointing to the possibility that a complete Om14 is required for optimal interaction with Mim1. The compromised organelle specificity in the targeting of the truncated constructs might be explained by the absence of a clear involvement of Mim1. Of note, a construct containing the first two TMDs (GFP-Om14(1-95)) was targeted to mitochondria in a higher extent as compared to the constructs harboring single TMDs (Fig. 1B and C). Collectively, we conclude that each one of the putative TMDs and its flanking region is sufficient for mitochondrial targeting, although with compromised specificity. An improved specificity is achieved with a protein fragment with two TMDs but, only the combination of all three segments in one protein assures the high mitochondrial specificity observed for native Om14.

### Tom70 plays only a minor role in the biogenesis of Om14

Obtaining this insight on the mitochondrial targeting information, we next aimed to understand how these signals are recognized at the organelle surface. An obvious candidate to function as an import receptor for Om14 is Tom70 that was reported to contribute to the biogenesis of multi-span MOM proteins in both yeast and human cells (Papić et al., 2011; Becker et al., 2011; Otera et al., 2007). To investigate the involvement of Tom70 in the biogenesis of Om14, we first characterized the interaction interface between Tom70 and Om14 using a peptide scan blot assay. Peptides of 20 amino acid residues with a shift of three amino acids covering the whole sequence of Om14 were synthesized as spots on a cellulose membrane. This membrane was incubated with GST-tagged cytosolic domain of Tom70 expressed recombinantly in *E. coli* cells. We initially verified that GST alone does not bind in this assay (data not shown). Using GST-Tom70, we observed that the strongest interaction occurred to peptides representing the C-terminal region of Om14 whereas moderate binding was detected with peptides residing inside or close to the first two predicted transmembrane domains of the protein (Fig. 2A). These findings clearly indicate that Tom70 does not bind to only a single well-defined segment of Om14.

**Figure 2.**
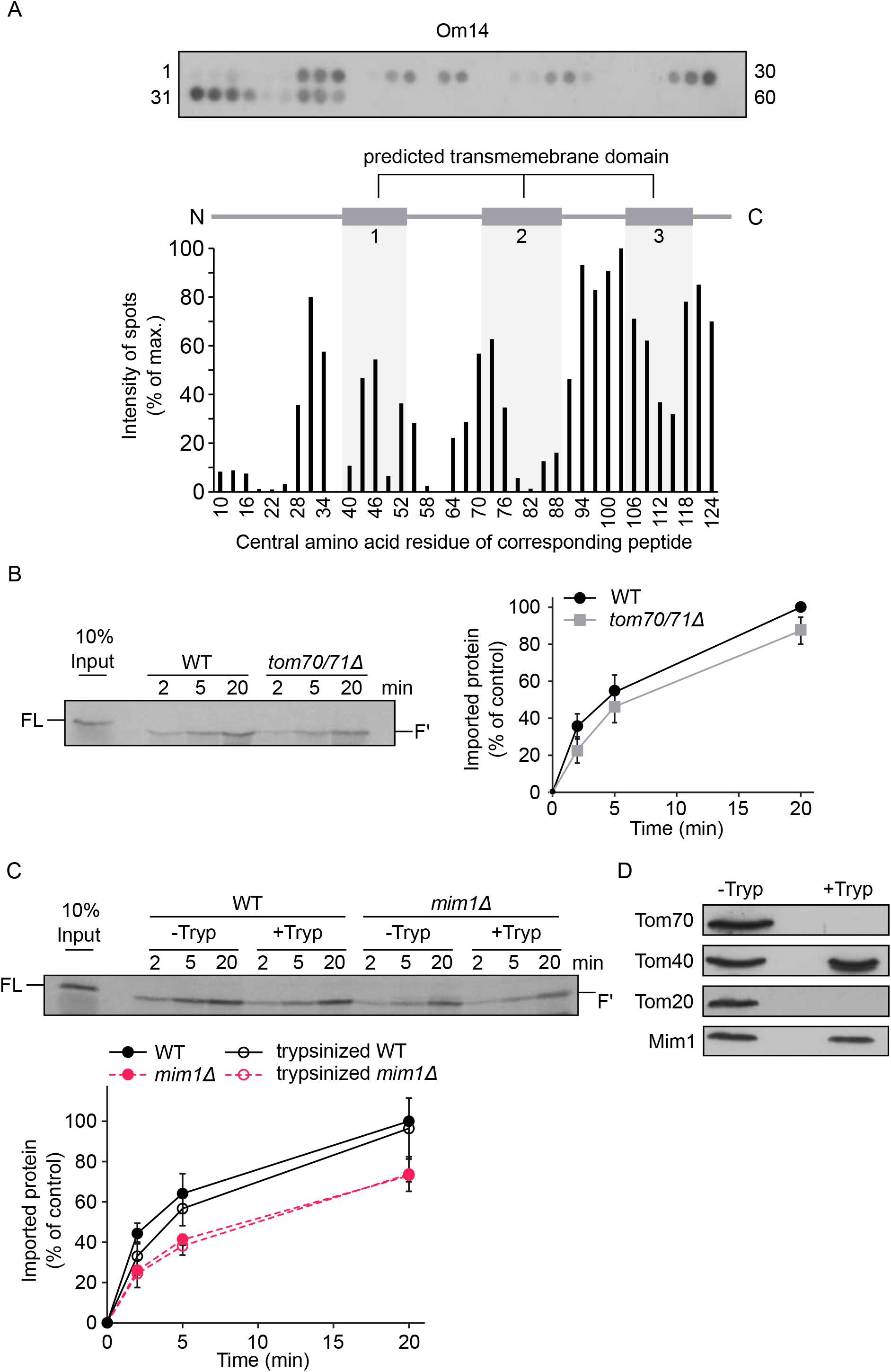
Loss of Tom70 has a minor impact on the biogenesis of Om14. **(A)** Nitrocellulose membrane containing 20-mer peptides covering the sequence of Om14 was incubated with the recombinant fusion protein GST-Tom70(cytosolic domain). After incubation, the interaction was visualized by immunodecoration using antibody against GST (upper panel). The numbers flanking the panel reflect the serial numbers of the peptides. Lower panel: The intensity of each dot was quantified and the average quantification of three independent experiments were plotted. The dot with the strongest intensity was set as 100%. The numbers on the X-axis reflect the central amino acid residue of each peptide. **(B)** Radiolabeled Om14 was imported into mitochondria isolated from either WT or *tom70/71Δ* cells. After import, samples were treated with trypsin and were further analyzed by SDS-PAGE and autoradiography. FL, full length input; F’, the trypsin-related proteolytic fragment of Om14. Right panel: The intensity of the trypsin-related fragment (F’) was quantified and the average intensity of three independent experiments is shown. The band representing import for 20 min into control organelles was set to 100%. **(C)** Mitochondria isolated from either WT or *mim1Δ* cells were left intact (-Tryp) or were pre-treated with trypsin (+Tryp). Radiolabeled Om14 was then added to the isolated organelles for the indicated time periods. After import, samples were again trypsinized and analyzed by SDS-PAGE and autoradiography. Lower panel: Quantification of three independent experiments was performed as described in (B). **(D)** Mitochondria isolated from WT cells were either left intact or treated with trypsin. Both samples were analyzed by SDS- PAGE and immunodecoration using the indicated antibodies.

To elaborate the physiological relevance of Tom70 for the biogenesis of Om14, we conducted an *in vitro* import assay in which newly synthesized radiolabeled Om14 was incubated with mitochondria isolated from either WT cells or from cells of a strain where *TOM70* and its paralogue *TOM71* were deleted (*tom70/71Δ*). As a read-out for the correct membrane integration of Om14, we utilized the formation of a characteristic 13 kDa fragment (F’) upon addition of external trypsin (Fig. S2A and Burri et al., 2006). We observed a reduction of about 10-15% in the integration capacity of Om14 into organelles lacking both Tom70 and Tom71 (Fig. 2B). We therefore conclude that while Tom70/71 contributes to the biogenesis of Om14, a large portion of newly synthesized Om14 molecules can be integrated into the MOM also in the absence of Tom70/71.

To study whether the involvement of Tom70 is evolutionary conserved, we performed the same import assay using mitochondria isolated from human cells depleted of TOM70. Interestingly, although no homolog of Om14 was found in the human genome, Om14 was still able to be inserted into mammalian organelles in a correct topology (Fig. S2B). Compared to yeast mitochondria, Om14 showed a slightly higher dependence on Tom70 in human mitochondria as reflected by a 30% import reduction upon knocking down TOM70 (Fig. S2B). Thus, in agreement with a previous report on the importance of TOM70 for the biogenesis of mammalian multi-span MOM proteins (Otera et al., 2007), Om14 can be integrated into mammalian MOM in a TOM70-dependent manner.

The rather minor import reduction of Om14 in the absence of Tom70/71 might suggest that other cytosol exposed proteins like Tom20 can take over the receptor function in the absence of Tom70/71. To abolish simultaneously the receptor function of Tom20 and Tom70, we pre-treated isolated mitochondria with trypsin prior to the import reaction. Although this treatment completely removed the cytosolic domains of both receptors (Fig. 2D), we observed only a minor reduction in Om14 import into mitochondria treated in this way (Fig 2C). Hence, it appears that both Tom70 and Tom20, or any other MOM proteins exposed to the cytosol, play only a subordinate role in the biogenesis of Om14. Of note, Mim1 was not affected by the addition of trypsin (Fig. 2D). Hence, we next wanted to investigate the involvement of Mim1 in the membrane integration of Om14. In line with a previous publication (Becker et al., 2011), the import of Om14 into *mim1Δ* mitochondria was significantly impaired (Fig. 2D). However, even in the absence of Mim1, the removal of exposed proteins by pre-treatment with trypsin did not cause further reduction in the import capacity of these mitochondria (Fig. 2D). Hence, it seems that the general import receptors Tom20 and Tom70/71 do not contribute to the remaining 60% import capacity in the absence of Mim1.

Next, we asked whether the cytosolic domain of Mim1 might function as a receptor for Om14 substrate molecules. To that end, we introduced into *mim1Δ* cells plasmid-encoded native Mim1 or a variant where only the central TMD of Mim1 (residues 35-75, Mim1-TMD) was present thus, eliminating the putative cytosol-exposed region. We previously demonstrated that over-expression of this variant can rescue the growth phenotype in the absence of native Mim1 (Popov-Čeleketić et al., 2008). Importantly, the presence of this truncated construct could rescue the steady-state levels of known MIM1 substrates, including Om14, to the same extent as native Mim1 (Fig. S3A). Furthermore, organelles harboring this variant could facilitate assembly of newly synthesized Om14 molecules as good as native Mim1 (Fig. S3B). Collectively, these results demonstrate that Tom70, Tom20, or the cytosolic region of Mim1 do not play a decisive role in the biogenesis of Om14.

### Hydrophobicity of the second TMD influences the membrane integration capacity of Om14

Whereas the membrane integration of Om14 is clearly promoted by Mim1, we wondered whether we could identify cis elements that affect this process. Thus, we had a closer look at the hydrophobicity of the three putative TMDs. Using Wimley and White hydrophobicity scale (Wimley and White, 1996), we noticed that the first TMD of Om14 displays the lowest hydrophobicity (1.0) whereas the other two TMDs have a similar value of 1.5. To determine how the hydrophobicity of TMDs affects the biogenesis of Om14, we analyzed the membrane integration capacity of two constructs that exhibit either an increased hydrophobicity of the first TMD (Om14-3I) or a decreased hydrophobicity of the second TMD (Om14-4A) (Fig. S4A and Fig. 3A, respectively). Om14-3I showed a similar topology and membrane integration capacity as the native protein (Fig. S4B), suggesting that the hydrophobicity of the first TMD does not affect the overall biogenesis of the protein. In contrast, replacing four hydrophobic residues of TMD2 by alanine, as in Om14-4A, caused a dramatic reduction in both the steady-state levels and the integration capacity of the protein (Fig. 3B and C). Notably, although Om14 contains three putative transmembrane segments, previous reports and our current experiments find a sub-population of Om14 in the supernatant fraction (Fig. 3B, Burri et al., 2006; Sauerwald et al., 2015). About 70% of the OM14-4A molecules, as compared to ca. 30% of the native protein, were found in the supernatant fraction. This altered membrane integration points to the possibility that the mutated variant is less stable and accordingly, the reduced steady-state levels might result from enhanced turn-over.

**Figure 3.**
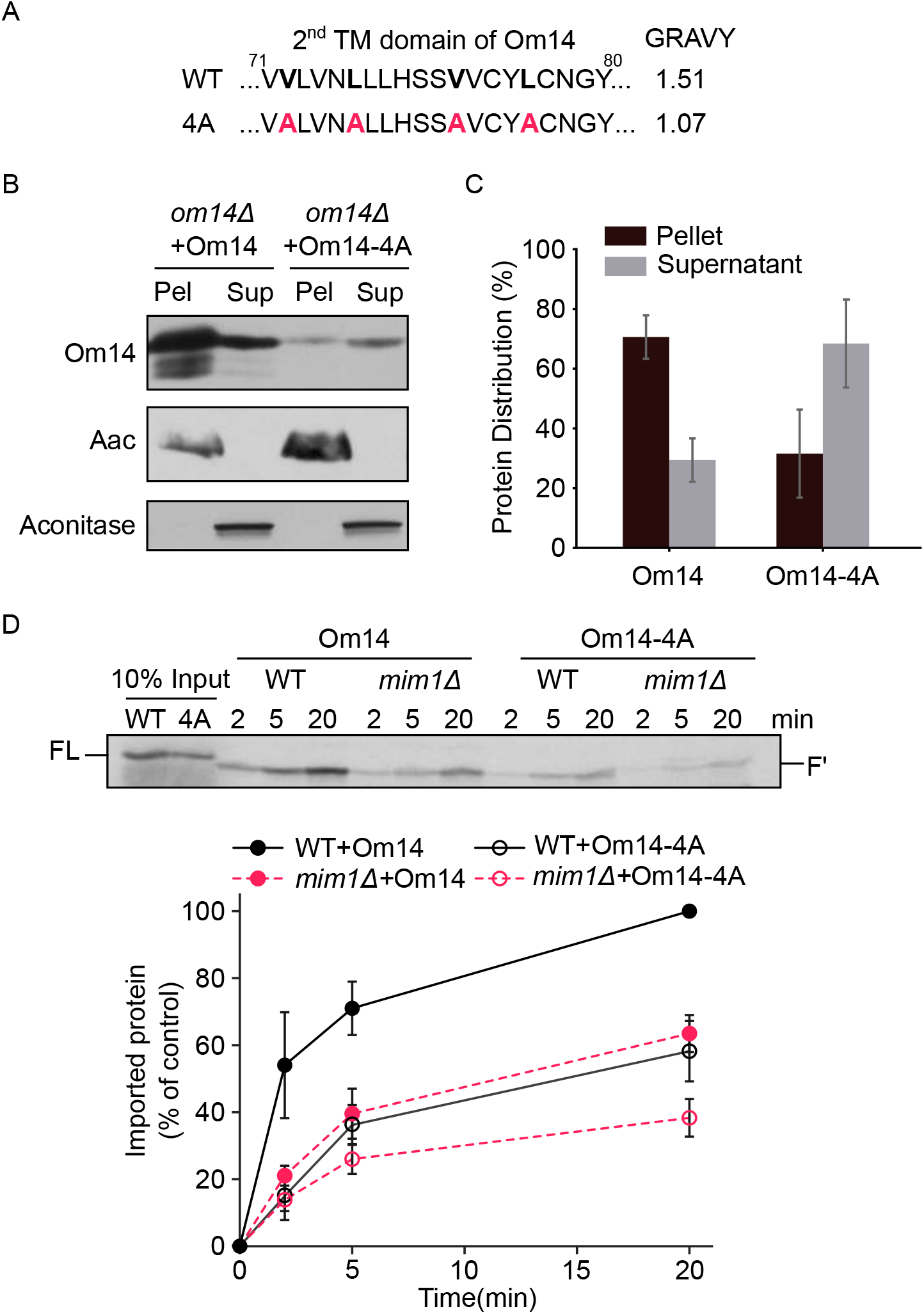
Reduced hydrophobicity of TMD2 compromises the import capacity of Om14. **(A)** Amino acid sequence of the putative second TMD of both native Om14 (WT) and a variant with reduced hydrophobicity due to the introduction of four Ala residues (4A). Mutated residues are shown in bold. **(B)** Mitochondria isolated from *om14Δ* yeast strains expressing either native Om14 or Om14-4A were subjected to alkaline extraction. Pellet (P) and supernatant (S) fractions were analyzed by SDS-PAGE and immunodecoration with antibodies against the indicated proteins. ATP-ADP carrier (Aac) is embedded in the mitochondrial inner membrane whereas Aconitase is a matrix soluble protein. **(C)** Quantification of at least three independent experiments as in (B). The combined intensity of Om14 in the pellet and supernatant fractions from each strain was set to 100%. **(D)** Radiolabeled versions of Om14 and its Om14-4A variant were imported into mitochondria isolated from either WT or *mim1Δ* cells. At the end of the import reactions, mitochondria were treated with trypsin and further analysis and quantification were as described in the legend to Fig. 2B. Lower panel: Quantification of three independent experiments is presented. The intensity of the band corresponding to import of native Om14 into control organelles for 20 min was set to 100%.

To address the question which stage of Om14-4A biogenesis was compromised and whether the obstruction was Mim1-dependent, we imported *in vitro* radiolabeled Om14-4A into organelles isolated from either control or *mim1Δ* cells. As revealed in Fig. 3D, the *in vitro* import efficiency of Om14-4A was largely compromised compared to native Om14. The import of Om14-4A was further impaired in the absence of Mim1, suggesting that the loss of integration capacity was probably not triggered by a reduced interaction of the variant with Mim1. Taken together, these results support the assumption that reduced hydrophobicity of TMD2 of Om14 interferes with MIM complex-independent membrane integration steps.

### Unfolding and elevated temperature strongly accelerate the import of Om14

Our results so far indicate that the complete length of the protein is required for optimal biogenesis of Om14. This led us to ask whether partial folding events are required to support the import of Om14. To address this point, we denatured *in vitro* translated Om14 using 6 M urea prior to the import reaction. Interestingly, this treatment enhanced four-fold the import capacity into control organelles (Fig. 4A). Of note, such an increase was observed also upon import into mitochondria lacking Mim1 (Fig. 4A), suggesting that the enhanced capacity upon unfolding is not related to a better interaction of the unfolded substrates with the MIM complex. These observations suggest that an optimal biogenesis of Om14 is favored by its unfolding in the initial stages arguing against the requirement of a tertiary structural targeting signal. Considering that many of the effects that we detected so far do not necessarily depend on a proteinaceous element, we wondered whether increasing the fluidity of the membrane by elevated temperature can also improve the biogenesis of Om14. To test this issue, we performed the *in vitro* import reaction while incubating the isolated organelles at either 25°C or 37°C. Importantly, we observed an increased import efficiency of Om14 after incubation at 37°C (Fig. 4B). To test whether this behavior is shared by other proteins, we also imported *in vitro* other MOM proteins like Ugo1 or Porin under the same conditions. As shown in Fig. 4C and D, the elevated temperature resulted in an increase of the import efficiency of Ugo1, although only after 20 min. In contrast, the elevated temperature resulted in a reduced import of the β-barrel protein Porin. Taken together, although it seems that increased fluidity of the membrane supports enhanced membrane integration of Om14, we cannot exclude the possibility that the beneficial effect of the higher temperature is also partially due to enhanced unfolding of the Om14 substrate.

**Figure 4.**
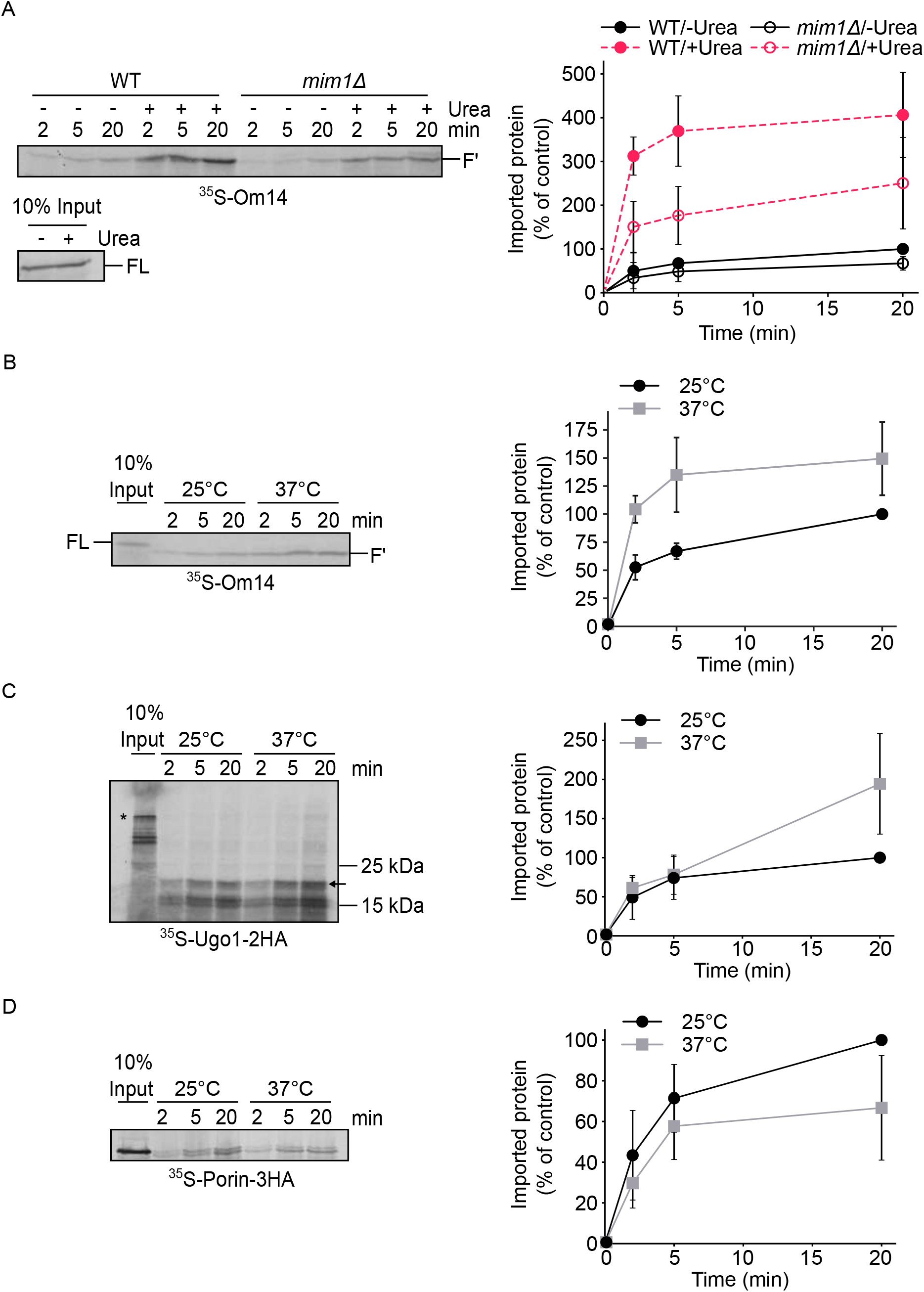
Unfolding and elevated temperature facilitate the import of Om14. **(A)** Radiolabeled Om14 was either denaturated in 6 M urea or left untreated before its import into mitochondria isolated from either WT or *mim1Δ* strains. After the import, mitochondria were treated with trypsin and further analysis and quantification were done as described in the legend to Fig. 2B. Right panel: Quantification of three independent experiments is presented. The intensity of the band corresponding to import of native Om14 into control organelles for 20 min was set to 100%. **(B-D)** Radiolabeled Om14 (B), Ugo1-2HA (C), or Porin-3HA (D) were imported at either 25°C or 37°C into mitochondria isolated from WT cells. At the end of the import reactions, mitochondria were treated with either trypsin (B, C) or PK (D) and further analyzed by SDS-PAGE and autoradiography. Right panels: quantification of the typical proteolytic fragments of Om14 (B) or Ugo1 (C) as well as the PK-protected correctly inserted Porin (D) of three independent experiments are shown. The intensity of the band corresponding to import for 20 min at 25°C was set to 100%.

### Porin and Om45 regulate different stages in the biogenesis of Om14

The aforementioned findings imply that mitochondria lacking Mim1, Tom20, and Tom70 can still maintain more than half of their capacity to import Om14. Hence, it seems that additional proteins might be involved in this process. As interaction partners of Om14, Porin and Om45 might be such mediators. It was previously reported that the steady-state level of Om14 was not notably affected in *om45Δ* mitochondria whereas deletion of Porin resulted in a reduction in Om14 levels (Lauffer et al., 2012). To better understand the dependence of Om14 on Porin and Om45, we isolated mitochondria from WT and from cells lacking either Porin or Om45 and separated peripheral protein from integral protein by alkaline extraction. We observed a drastic increase of Om14 in the supernatant fraction of *om45Δ* cells whereas the absence of Porin did not affect the behavior of Om14 (Fig. 5A, B). These findings indicate an altered membrane association of Om14 upon the loss of Om45. Furthermore, the absence of Om45 also caused a significant shift in the migration of Om14-containing oligomeric structures as observed by BN-PAGE (Fig. 5C). As a further approach, we imported *in vitro* radiolabeled Om14 into mitochondria isolated from WT, *om45Δ,* or *por1Δ* strains and analyzed membrane integration of Om14 with the trypsinization assay. The import efficiency of Om14 dropped by about 20% or 30% in mitochondria lacking either Om45 or Porin, respectively (Fig. 5D). These findings indicated that both Porin and Om45 contribute to the membrane integration of Om14 whereas Om45 seems to play a more important role in the downstream stages of complex assembly. Since a similar import reduction was observed in the absence of either Mim1 or Porin (compared Fig. 2C to Fig. 5D), we speculated that Mim1 and Porin might compensate for the loss of each other in mediating the early biogenesis of Om14. To test this possibility, we generated a *por1Δmim1Δ* double deletion mutant and analyzed the steady state level of Om14 in these cells. We observed that the level of Om14 in the double deletion mutant was drastically reduced while the reduction fits the additive effect of the single deletions (Fig. 6A and B). Importantly, the steady-state levels of other MIM substrates like Tom70 and Tom20 were reduced upon the deletion of Mim1 but were not significantly further reduced upon the co-deletion of Porin (Fig. 6A and B). This observation illustrates the specificity in the contribution of Porin to the biogenesis of Om14.

**Figure 5.**
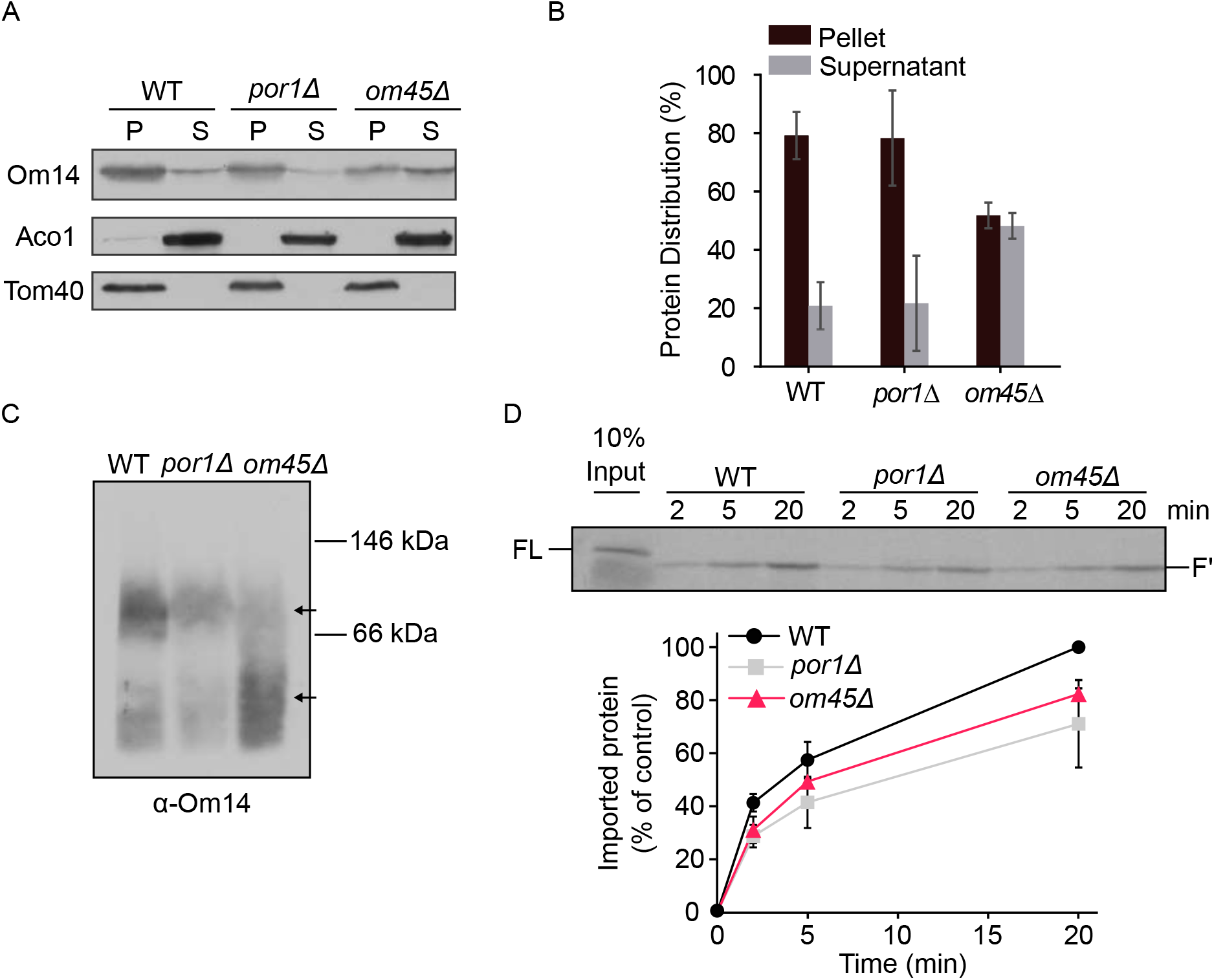
Om45 regulates the stability of Om14 oligomeric structures. **(A)** Mitochondria isolated from the indicated cells were subjected to alkaline extraction. Pellet (P) and supernatant (S) fractions were analyzed by SDS-PAGE and immunodecoration with the indicated antibodies. Aconitase (Aco1) serves as a marker for soluble matrix proteins whereas Tom40 is embedded in the mitochondrial outer membrane. **(B)** Quantification of three independent experiments as in (A). The combined intensity of Om14 in the pellet and supernatant fractions from each cell type was set to 100%. Error bars represent ±SD. **(C)** Mitochondria isolated from WT, *por1Δ*, or *om45Δ* cells were lysed using digitonin and analyzed by BN-PAGE. The blot was immunodecorated with an antibody against Om14. Arrows indicate the migration of oligomeric species of Om14. **(D)** Radiolabeled Om14 was imported into mitochondria isolated from WT, *por1Δ*, or *om45Δ* cells. at the end of the import reactions, mitochondria were treated with trypsin and analyzed by SDS-PAGE and autoradiography. Lower panel: The intensity of the proteolytic fragment (F’) in three independent experiments was quantified as described in the legend to Fig. 1B. The intensity of the band corresponding to import into control organelles for 20 min was set to 100%.

**Figure 6.**
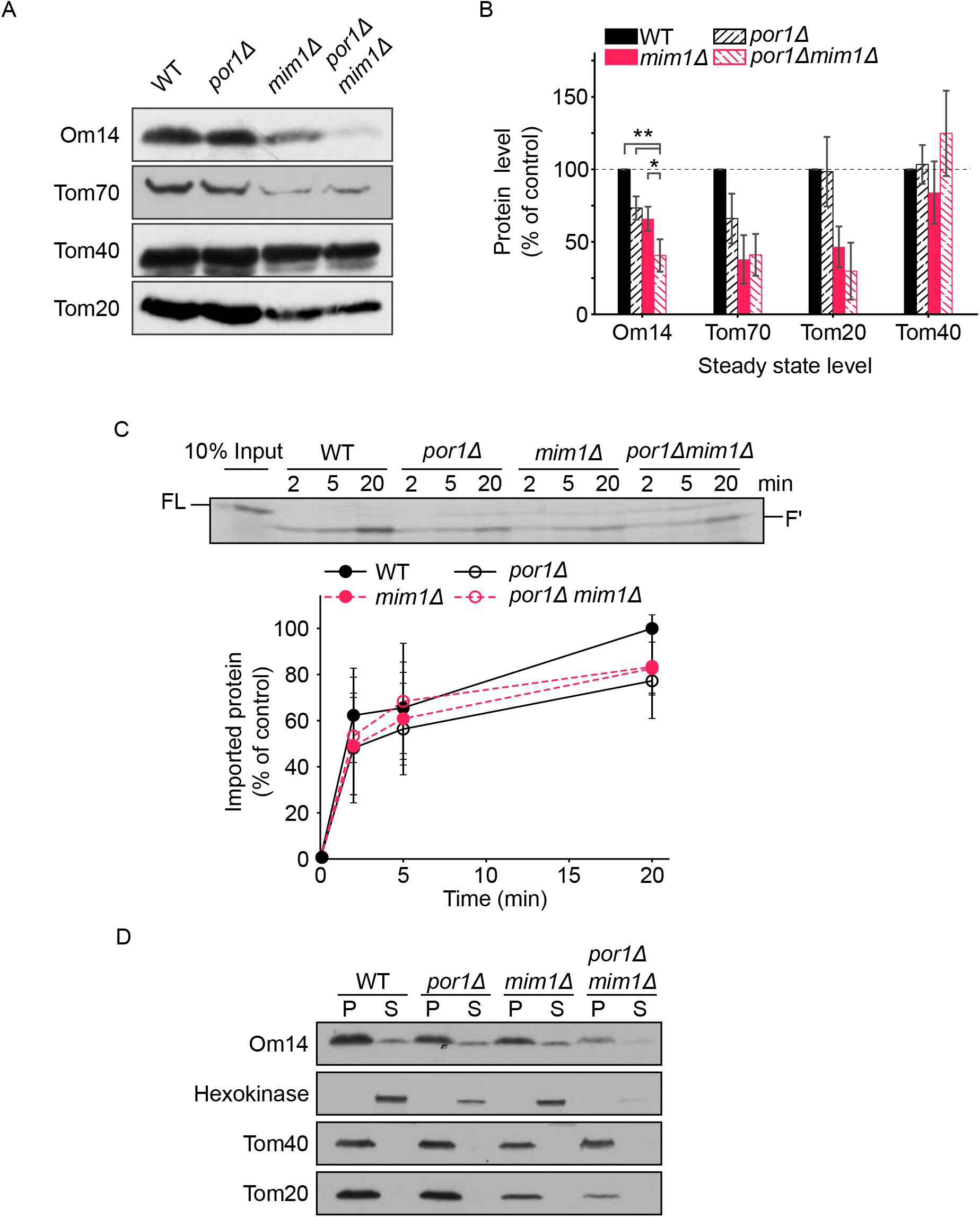
Both Mim1 and Porin contribute to the biogenesis of Om14. **(A)** Mitochondria isolated from WT, *por1Δ*, *mim1Δ*, or *por1Δmim1Δ* cells were analyzed by SDS-PAGE and immunodecoration with antibodies against the indicated proteins. **(B)** The steady-state levels of the proteins analyzed in (A) were quantified and the levels in the WT cells were set to 100%. Error bars represent ±SD. n>3; *, p≤0.05; **, p≤0.01. **(C)** Radiolabeled Om14 was imported into mitochondria isolated from the cells described in (A). At the end of the import reactions, mitochondria were treated with trypsin and the import reactions were further analyzed and quantified as described in the legend to Fig. 2B. Lower panel: Quantification of three independent experiments is presented. The intensity of the band corresponding to import into control organelles for 20 min was set to 100%. **(D)** Mitochondria as in (A) were subjected to alkaline extraction. Pellet (P) and supernatant (S) fractions were analyzed by SDS-PAGE and immunodecoration using the indicated antibodies. A sub-population of Hexokinase is associated peripherally to the surface of mitochondria via Porin (Por1). Tom20 is anchored to the mitochondrial outer membrane.

To further test whether the reduction in the steady-state levels of Om14 in *por1Δmim1Δ* cells resulted from downstream effects like inefficient assembly or instability of integrated Om14, we imported *in vitro* radiolabeled Om14 into organelles isolated from *por1Δ, mim1Δ,* or *por1Δmim1Δ* cells. Compared to the impairment of import efficiency in the single *por1Δ* or *mim1Δ* deletions, no further reduction in the import capacity was detected upon the double deletion (Fig. 6C). Finally, we tested whether the absence of both Mim1 and Porin would influence the membrane integration of Om14. To that goal, we monitored the portion of Om14 molecules in the supernatant fraction of alkaline extraction and found that, although the steady-state levels of Om14 were dramatically reduced, the extracted fraction was not increased in the double deletion strain (Fig. 6D). These results suggest that Mim1 and Porin are separately involved in mediating the early biogenesis steps of Om14 but, in contrast to Om45, both proteins do not contribute to the stability of Om14 within the membrane.

## Discussion

In the current study, we dissected the biogenesis pathways of the MOM multi-span protein Om14. Initially, we asked which segment(s) provide the mitochondrial targeting information within this protein. It is known that mitochondrial targeting of such proteins is not mediated by a canonical N-terminal presequence but rather by elements that are part of the mature form of the protein. However, the exact signals of such proteins where not resolved yet. We observed that truncated variants are targeted to mitochondria but in addition also to extra-mitochondrial locations. In this context, it is interesting that the removal of only several amino acid residues at the C-terminal flanking region of the first TMD was sufficient to compromise the mitochondrial targeting of this construct. Collectively, these results strongly support a model in which only the additive contribution of several local signals assure specific mitochondrial targeting. Currently, it is still unclear which factors decode this information.

A clear candidate for such a function was the import receptor Tom70, which appears to have dual function, on the one hand it can recognize internal mitochondrial signals (Brix et al., 1999; Papić et al., 2011), and on the other hand it serves as a docking site for cytosolic (co)chaperones on the mitochondrial surface (Young et al., 2003; Opaliński et al., 2018; Backes et al., 2021). Once Om14 is synthesized in the cytosol, co-chaperones of the Hsp40 family together with Hsp70 chaperones associate with it (Jores et al., 2018). Although the precise physiological relevance of this association was not studied yet, one can speculate that the (co)chaperones shield the hydrophobic segments and help to avoid misfolding and/or premature aggregation in the cytosol. The reported association of Om14 with cytosolic chaperones can explain, at least partially, the reported contribution of Tom70 for the biogenesis of such proteins (Becker et al., 2011; Papić et al., 2011).

Indeed, deletion of *TOM70* in yeast or knock-down of the protein in mammalian cells was found in the current study and was reported before to cause a reduction in the import of MOM multi-span proteins (Brix et al., 1999; Papić et al., 2011; Becker et al., 2011; Otera et al., 2007; Wan et al., 2016). However, the absence of Tom70 results often, including the current study, in only a moderate decrease in the import efficiency of these substrates. Such a variable dependency on Tom70 might be related to the different association of substrates with cytosolic chaperones and/or to the capacity of the substrates to use alternative routes. In addition, in the case of Om14, which is a very abundant protein (Morgenstern et al., 2017), one can speculate that its minor dependence on Tom70 might be a mechanism to avoid a situation where a large portion of Tom70 molecules is busy processing Om14 while leaving a limited capacity for other substrates.

The mild effect of the absence of Tom70/71 suggests the existence of alternative pathways. Interestingly, Om14 showed a higher Tom70 dependence in human cells, maybe because it is not a natural substrate of these cells and thus is not an optimal substrate of alternative pathways. Considering that the current model proposes that Tom70 relays the substrate proteins towards the insertase, the MIM complex (Fig. 7, pathway I), alternative routes might include direct recognition of substrates by the MIM complex (Fig. 7, route IIIa). However, our previous observation that the central TMD of Mim1, which lacks the putative cytosolic part of Mim1, can compensate for the growth phenotype in the absence of the native protein (Popov-Čeleketić et al., 2008) and our current observation that this domain supports normal levels of MIM substrates argue against a receptor-like function of Mim1.

**Figure 7.**
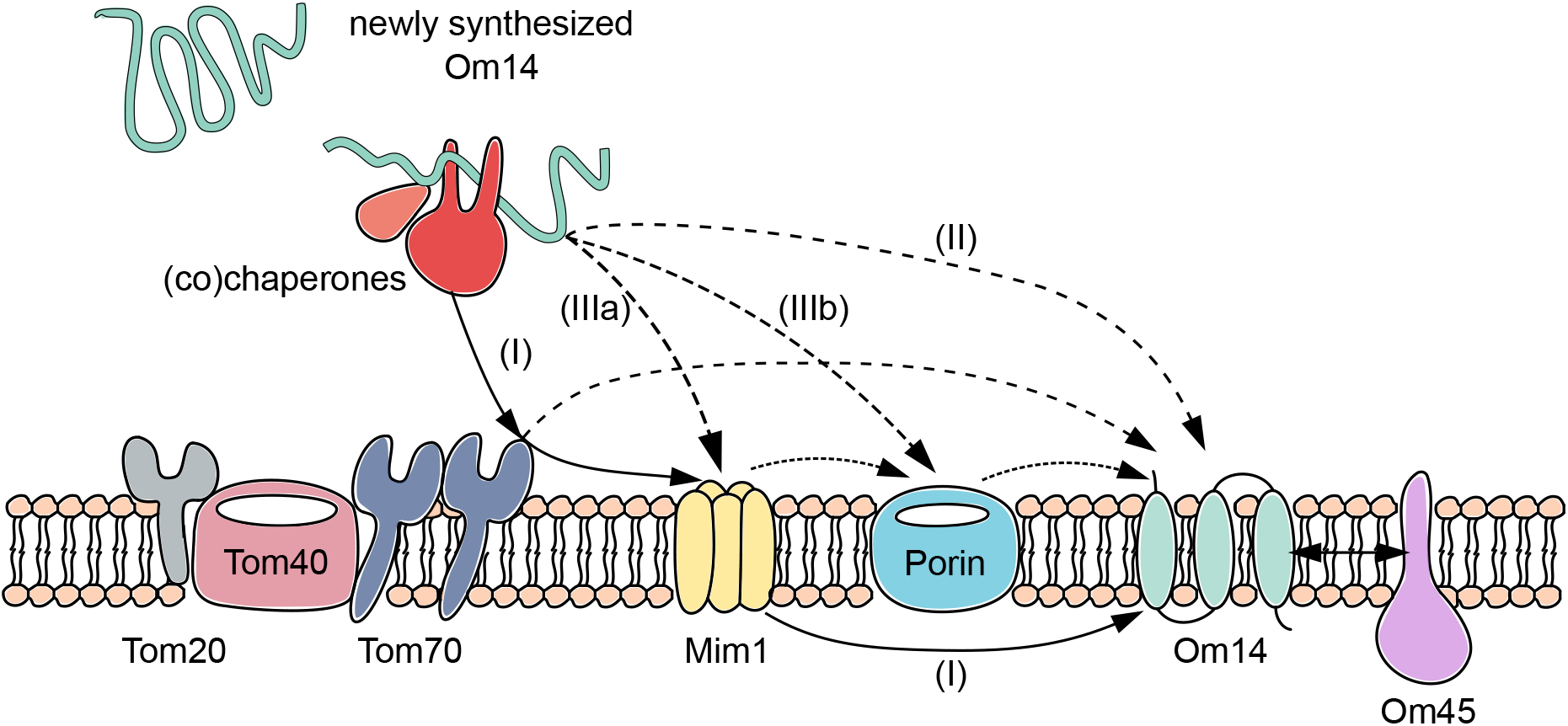
Various potential routes for the biogenesis of Om14. Newly synthesized molecules of Om14 can follow different pathways on their way to be integrated into the MOM. See the Discussion section of the main text for details.

A further additional option is direct recognition of Om14 substrate by Porin (Fig. 7, route IIIb), as our findings indicate a clear contribution of Porin to the correct membrane integration of Om14. However, Porin exposes only short loops towards the cytosol and therefore its function as a receptor is questionable. Hence, it seems that Porin promotes the correct membrane integration of Om14 but does not function as an initial receptor for the recognition of substrate molecules. This membrane-integration promoting function of Porin is yet another example for its multi-functionality in the biogenesis of other proteins. Recently, Porin was suggested to contribute to the assembly of the TOM complex by interacting with Tom22 (Sakaue et al., 2019) and to the import of carrier-like proteins to the mitochondrial inner membrane (Ellenrieder et al., 2019). The mechanism by which Porin can promote the membrane integration of Om14 should be the topic of future studies.

Importantly, even in the combined absence of both Mim1 and Porin we observed that a substantial portion of the Om14 molecules could still be properly integrated both *in vitro* and *in vivo* into the MOM. Hence it seems that either a yet unknown protein mediates this integration in the absence of these two proteins, or as suggested before by Vögtle et al. (2015), such multi-span proteins can be assembled into the membrane in an unassisted manner (Fig. 7, route II). Our findings that unfolding of Om14 or maintaining the hydrophobicity of its second TMD are important for the membrane integration of the protein support such direct substrate- lipids crosstalk. Further support for the importance of the membrane behavior is the increased membrane integration of Om14 at elevated temperature. As elevated temperature is known to enhance membrane fluidity (Kolodziej and Zammit, 1990), it is tempting to speculate that lowering the stiffness of the membrane increases the capacity of the hydrophobic segments of Om14 to integrate into the membrane. This suggestion is also in line with the previous observations of Vögtle et al. (2015) where the anionic phospholipid phosphatidic acid (PA), which is also known to induce curvature in membranes, supported membrane integration of the multi-span protein Ugo1. A further support for the increased dependency of Om14 integration on the membrane properties rather than on other proteinaceous factors is coming from the comparison to the reduced membrane integration of Porin at the higher temperature. The reduced import of this β-barrel protein reflects most likely its high dependency on the activity of multiple proteinaceous machineries like the TOM and TOB/SAM complexes.

Regardless of the precise route for the membrane integration of Om14, it appears that once the protein is embedded in the membrane, its interaction partner Om45 mediates its final maturation. The absence of Om45 resulted in enhanced fraction of Om14 molecules that can be extracted from the membrane as well as alteration in the oligomeric structures of Om14. Since in addition to Om45, also the hydrophobicity of the second TMD of Om14 appears to be important for its stable membrane integration, it might be that the second TMD of Om14 interacts with the putative N-terminally TMD of Om45.

Collectively, our findings provide a novel insight into the diversity of parameters affecting the biogenesis of a multi-span mitochondrial outer membrane protein. They reveal that rather than a single determinative factor, the import of Om14 is modulated by various membrane proteins and its own structural properties. Considering the outcome of a previous study suggesting the existence of multiple pathways regulating the biogenesis of MOM single-span proteins (Vitali et al., 2020), it seem that all the helical MOM proteins are integrated into the membrane in a process that is affected by cis elements in the substrate protein itself, a combination of trans proteinaceous elements, and the behavior of the membrane.

## Materials and methods

### Yeast strains and growth conditions

The *Saccharomyces cerevisiae* strains used in this study are listed in Table S1. The *por1Δmim1Δ* strain was created by mating the single deletion strains and then performing tetrad dissection. Yeast cells were grown at 30°C on selective media containing 2% of either galactose or lactate.

### Recombinant DNA techniques

Primers containing different restriction sites were designed to amplify by PCR segments encoding different regions of Om14. The plasmid pGEM4-Om14 was used as a template. The PCR products were subcloned into either the plasmid pGEM4 for *in vitro* protein translation or into the yeast expression vector pYX142-eGFP to study the *in vivo* behavior. The site-directed mutagenesis within Om14 was generated by using a pair of primers that contain mutated sequence of Om14 using pGEM4-Om14 as a template. The PCR products were digested using DpnI before being transformed into *E. coli* cells for further selection. Full lists of all primers and plasmids used in this study are found in Tables S2 an S3, respectively.

### Mitochondria isolation

Isolation of yeast mitochondria was performed by differential centrifugation following a previously published protocol (Daum et al., 1982). Isolated yeast mitochondria were stored at -80°C in SEM buffer (250 mM sucrose, 10 mM MOPS/KOH, 1 mM EDTA, pH 7.2). The mammalian HeLa cell line, which was used in this study, was a gift from Dr. Vera Kozjak-Pavlovic and doxycycline was used to induce knock-down of TOM70 as described previously (Kozjak-Pavlovic et al., 2007). For isolation of mammalian mitochondria, cells were washed once with PBS buffer and collected from the 150 mm tissue culture-treated dish using a spatula.

The collected cells were transferred to a test tube and were centrifuged (300*xg*, 5 min, 4°C). Afterwards, the cell pellet was resuspended in HMS buffer (0.22 M Mannitol, 0.02 M HEPES-KOH, pH 7.6, 1 mM EDTA, 0.07 M Sucrose, 0.1% BSA, 1 mM phenylmethylsulfonyl fluoride (PMSF)). The cells were then lysed nine times through each of the three different needles used (20G, 23G, and 27G, Sterican). After homogenization, samples were centrifuged (900*xg*, 5 min, 4°C) and the supernatant was centrifuged (9000*xg*, 15 min, 4°C). The pellet of the latter centrifugation step was washed with HMS buffer and re-isolated (10000*xg*, 15 min, 4°C). The pellet from the last centrifugation step represents the crude mitochondria and was used for further *in vitro* import experiment.

### *In vitro* protein import into mitochondria

Proteins encoded in pGEM4 plasmid were transcribed *in vitro* by SP6 polymerase. The acquired mRNA was then translated in rabbit reticulocyte lysates (Promega) containing ^35^S-labelled Met and Cys. After translation, ribosomes were removed (76000*xg,* 45 min, 4°C) and discarded. For an import reaction, mitochondria were diluted to a concentration of 1 µg/µl with F5 import buffer (250 mM sucrose, 0.25 mg/ml BSA, 80 mM KCl, 10 mM MOPS-KOH, 5 mM MgCl2, 8 mM ATP, 4 mM NADH, pH 7.2). In a standard import reaction, 10% (v/v) of translated protein was added into 50 µl of mitochondria solution and incubated at 25°C for 2, 5 or 20 min. In some experiments, mitochondria were incubated at 37°C in total for 20 min (before and during the import reaction). The import reactions were stopped by adding 400 µl of SEM-K^80^ buffer followed by a 10 min centrifugation (13200*xg*, 2°C). For further proteolytic assay, import reactions were treated for 25 min at 4°C with 50 µg/ml of either Proteinase K or Trypsin. The protease activity was terminated by incubation for 10 min at 4°C with either PMSF or trypsin inhibitor, respectively. Afterwards, samples were centrifuged (18000*xg* 10 min, 2°C) and the supernatants were discarded. For SDS-PAGE, the pellets were dissolved in 2x Sample buffer and then heated at 95°C for 10 min. Then, samples were loaded on either 12.5% or 15% SDS gels and blotted onto nitrocellulose membranes.

For analysis of import reactions with blue native (BN) PAGE, samples were dissolved in 1% (w/v) digitonin. After a clarifying spin (30000*xg*, 30 min, 2°C), the soluble supernatant was mixed with 10X BN loading dye. Samples were further analyzed by BN-PAGE and blotted onto PVDF membranes. The imported proteins were visualized by autoradiography and AIDA image analyzer was used to quantify the intensity of protein bands.

### Subcellular fractionation

Different cell fractions from yeast were separated using published subcellular fractionation protocol (Walther et al., 2009). To obtain pure mitochondria fraction, isolated mitochondria were slowly layered on top of a Percoll gradient for a better separation between mitochondria and contaminants (Graham, 2000). Proteins in whole cell lysate, cytosol, and ER fractions were precipitated by trichloroacetic acid (TCA) and finally dissolved in 2x Sample buffer and incubated at 95°C for 10 min before SDS-PAGE analysis. Proteins of interest were visualized by immnodecoration using the indicated antibodies. Table S4 indicates the antibodies used in the current study.

### Alkaline extraction

Isolated mitochondria (50 µg) were resuspended in 100 µl of 0.1 M Na2CO3 solution and incubated for 25 min at 4°C. Samples were centrifuged (76000*xg*, 30 min, 2°C), and the supernatant was collected and precipitated by TCA. Both pellets and precipitated proteins from the supernatant were dissolved in 40 µl of 2x Sample buffer and heated at 95°C for 10 min before further analysis by SDS-PAGE.

### Peptide scan assay

Cellulose membranes harboring 20-mer peptides covering the sequence of Om14 were activated in methanol for 1 min and washed twice with sterile water for 1 min. Then, membranes were equilibrated at room temperature with binding buffer (50 mM Tris-HCl, 150 mM NaCl, pH 7.5) for 2 hrs. After blocking with 1 µM BSA for 1 h, membranes were incubated overnight with 0.5 μM GST-Tom70-cytosolic domain. Next, the membranes were washed three times with binding buffer and bound protein was visualized by immunodecoration using antibody against GST.

### Fluorescence microscopy

Yeast cultures were initially grown overnight in synthetic media with 2% galactose. Cultures were then diluted to OD600 of 0.2 and after further growth, cells were harvested at the logarithmic phase (3000*xg*, 5 min, RT). The collected cells were mixed with 1% (w/v) low melting point agarose in a 1:1 (v/v) ratio and were plated on a glass slide. Spinning disk microscope Zeiss Axio Examiner Z1 equipped with a CSU-X1 real-time confocal system (Visitron) and SPOT Flex CCD camera (Visitron Systems) was used and Images were acquired using the AxioVision software (Visitron).

## Acknowledgments

We thank E. Kracker for excellent technical assistance, K. Maruszczak and Elias Schermuly for help with some experiments, V. Kozjak-Pavlovic for cells, and J. Nunnari and T. Becker for antibodies. This work was supported by the Deutsche Forschungsgemeinschaft (RA 1028/7-2 and 10-2 to D.R. and CRC 894 to M.J.).

## Author contributions

J.Z. and K.S.D. designed and conducted experiments, M.J. prepared the peptide scan membranes, D.R. designed experiments and analyzed data, J.Z. and D.R. wrote the manuscript.

---

**Table S1.** Yeast strains used in this study.

| Name | Mating type | Genetic background | Source of reference |
| --- | --- | --- | --- |
| W303a | MATa | *ade2-1 can1-1100 his3-11 leu2 3_112 trp1Δ2 ura3-52* | Lab stock |
| W303α | MATα | *ade2-1 can1-1100 his3-11 leu2 3_112 trp1Δ2 ura3-52* | Lab stock |
| *mim1Δ* | MATa | W303a; *mim1Δ::KanMX* | (Dimmer et al. 2012) |
| *mim1Δ* | MATα | W303α; *mim1Δ::KanMX* | (Dimmer et al. 2012) |
| *tom70Δ/71Δ* | MATα | W303α; *tom70Δ::KanMX tom71Δ::Nat* | Lab stock |
| *por1Δ* | MATα | W303α; *por1Δ::His3* | Lab stock |
| *om45Δ* | MATα | W303α; *om45Δ::His3* | Lab stock |
| *om14Δ* | MATa | W303a; *om14Δ::His3* | Lab stock |
| *por1Δ /mim1Δ* | MATa | W303α; *por1Δ::His3 mim1Δ::KanMX* | This study |

**Table S2.**
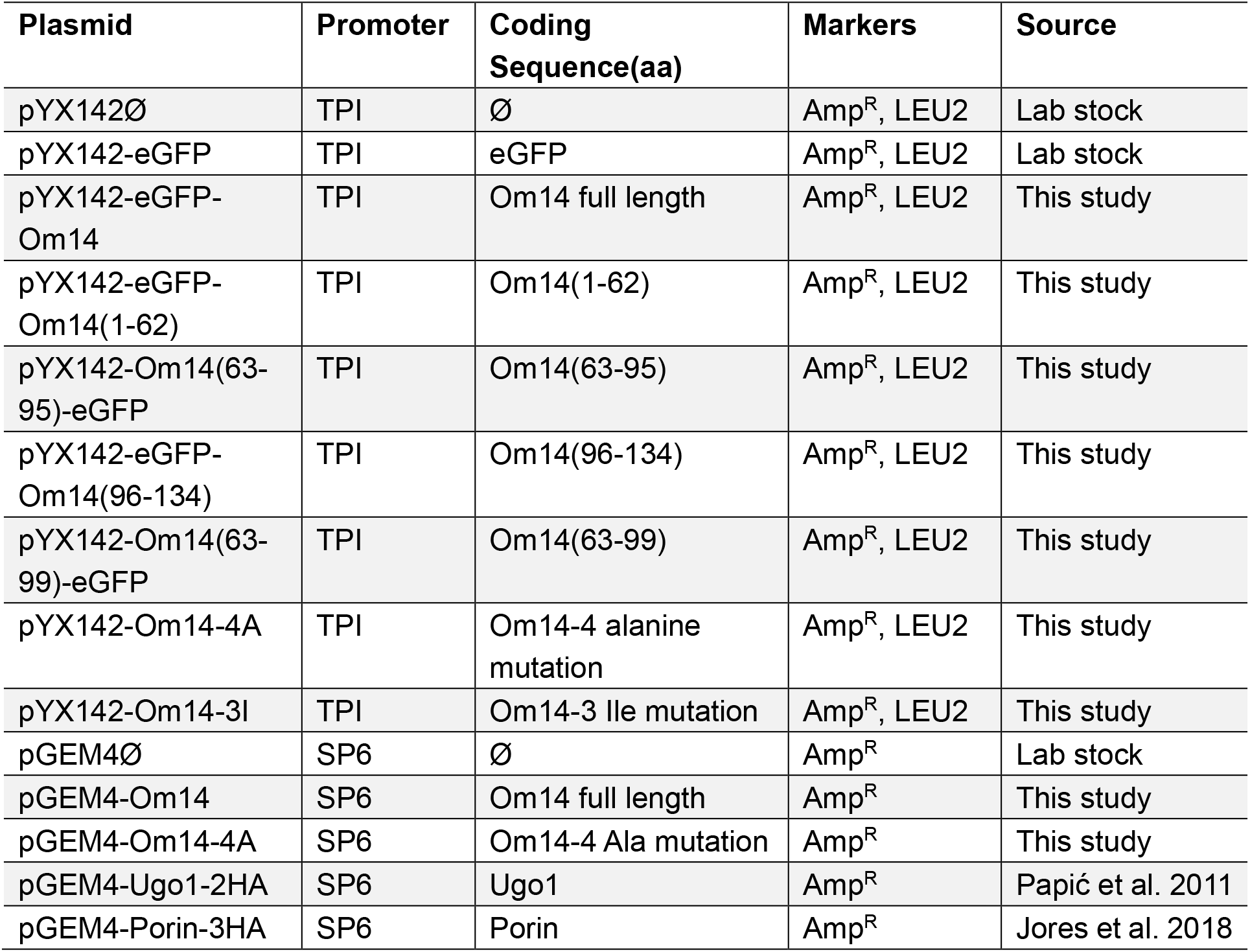
Plasmids used in this study.

**Table S3.**
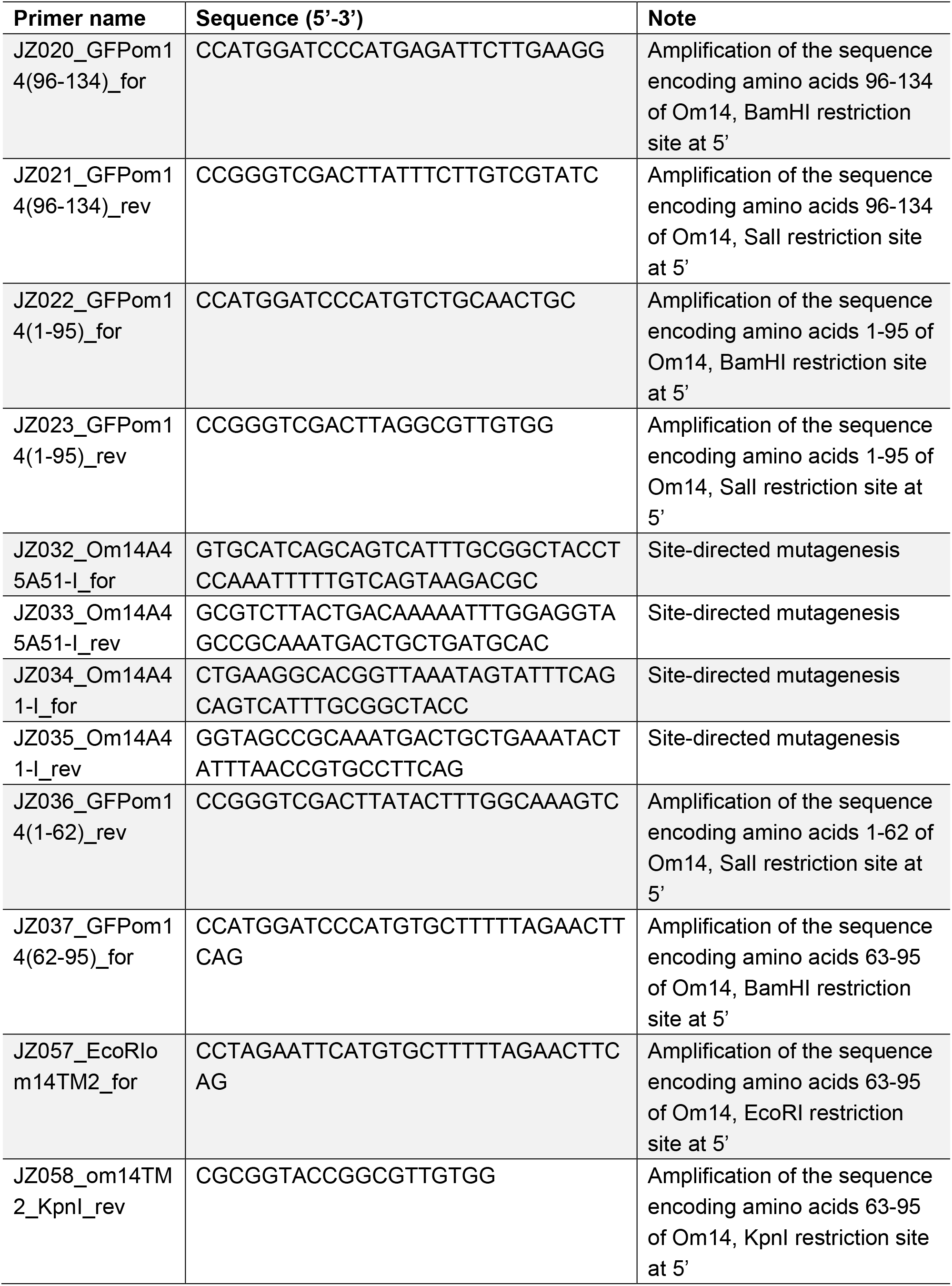
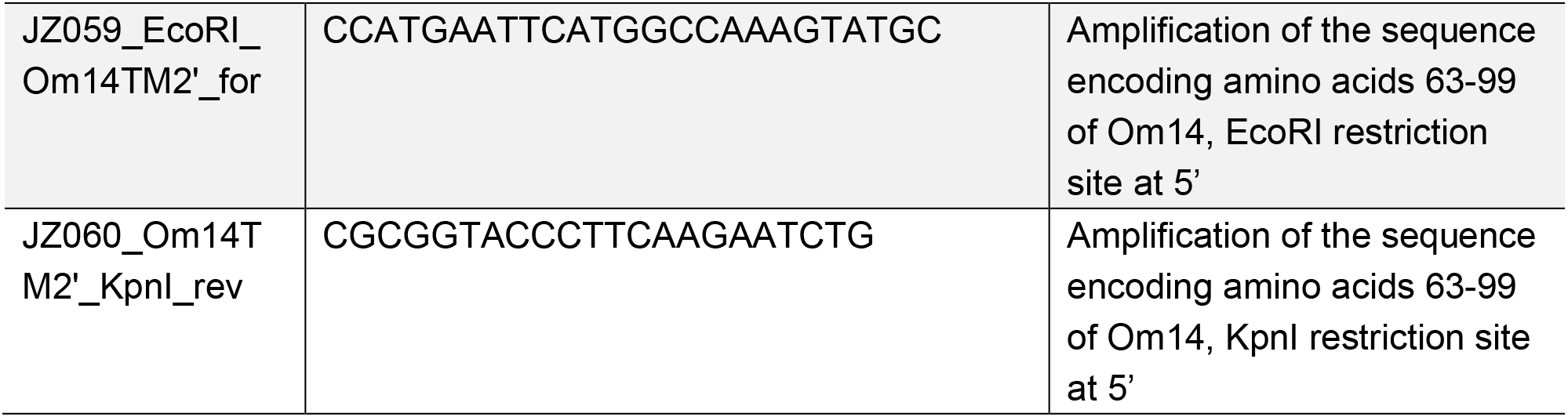
Primers used in this study.

**Table S4.** Antibodies used in this study.

| **Antibodies** | **Dilution** | **Source** |
| --- | --- | --- |
| polyclonal rabbit anti-GFP | 1:2000 | TP401 (Torrey Pines) |
| polyclonal rabbit anti-Tom70 | 1:2000 | Lab stocks |
| polyclonal rabbit anti-Tom40 | 1:5000 | Lab stocks |
| polyclonal rabbit anti-Tom20 | 1:2000 | Lab stocks |
| polyclonal rabbit anti-Mim1 | 1:500 | Lab stocks |
| polyclonal rabbit anti-Om14 | 1:2000 | Lab of Thomas Becker |
| polyclonal rabbit anti-Aco1 | 1:10000 | Lab stocks |
| polyclonal rabbit anti-Hexokinase | 1:3000 | Bio-Trend (#100-4159) |
| polyclonal mouse anti-AAC | 1:5000 | Lab stocks |
| polyclonal rabbit anti-Erv2 | 1:2000 | Lab stocks |
| polyclonal rabbit anti-Scm4 | 1:250 | Lab of Thomas Becker |
| polyclonal rabbit anti-Porin | 1:3000 | Lab stocks |
| polyclonal rabbit anti-Ugo1 | 1:2000 | Lab of Jodi Nunnari |

**Figure S1.**
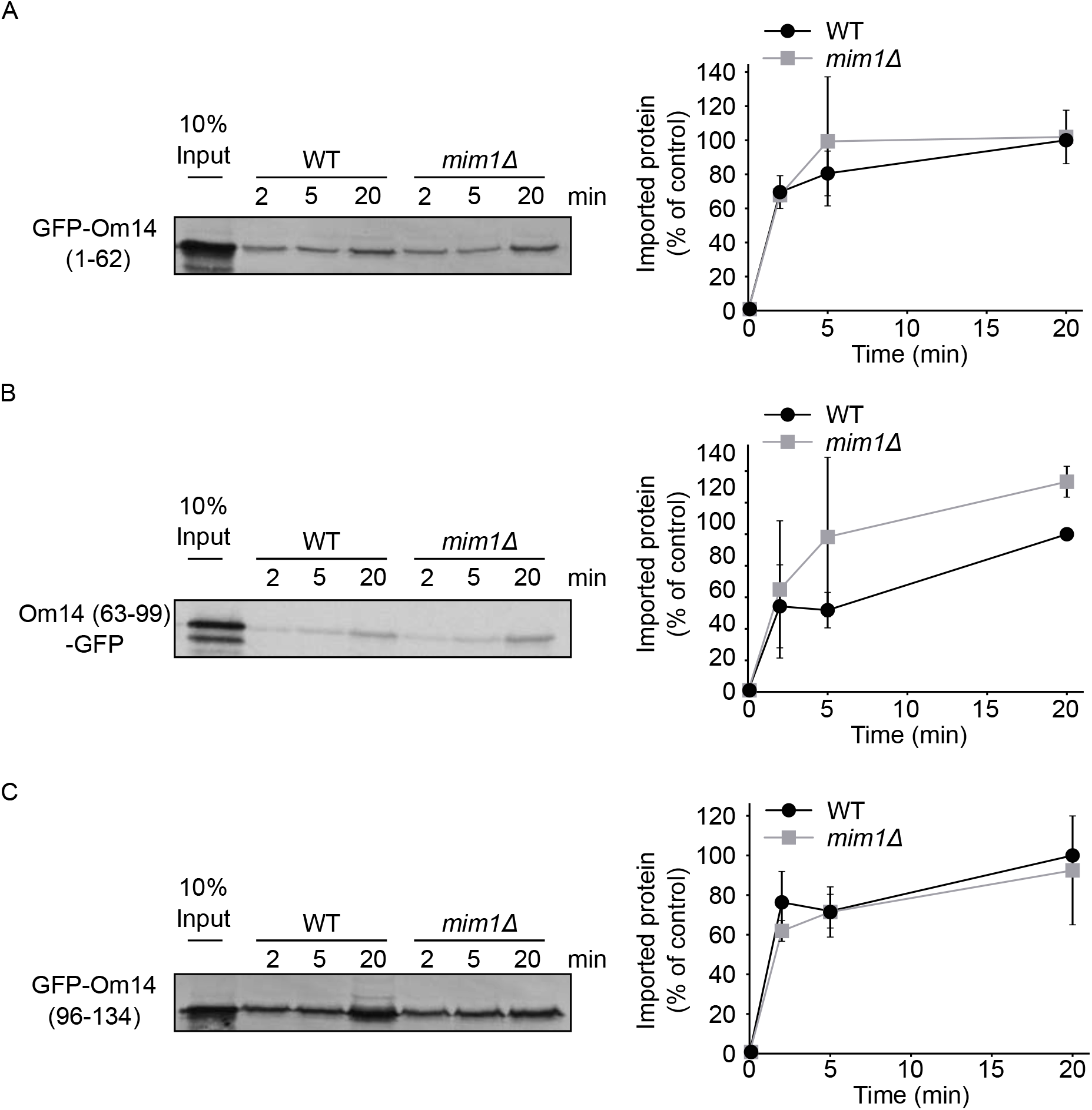
The import of truncated Om14 variants is only mildly dependent on Mim1. (A-C) Radiolabeled truncated variants of Om14 fused to GFP were incubated with mitochondria isolated from either WT or *mim1Δ* cells. At the end of the import reactions, samples were subjected to alkaline extraction and the pellets fractions were analyzed by SDS-PAGE and autoradiography. Right panels: Quantification of three independent experiments. The intensity of the band corresponding to import for 20 min into control organelles was set to 100%.

**Figure S2.**
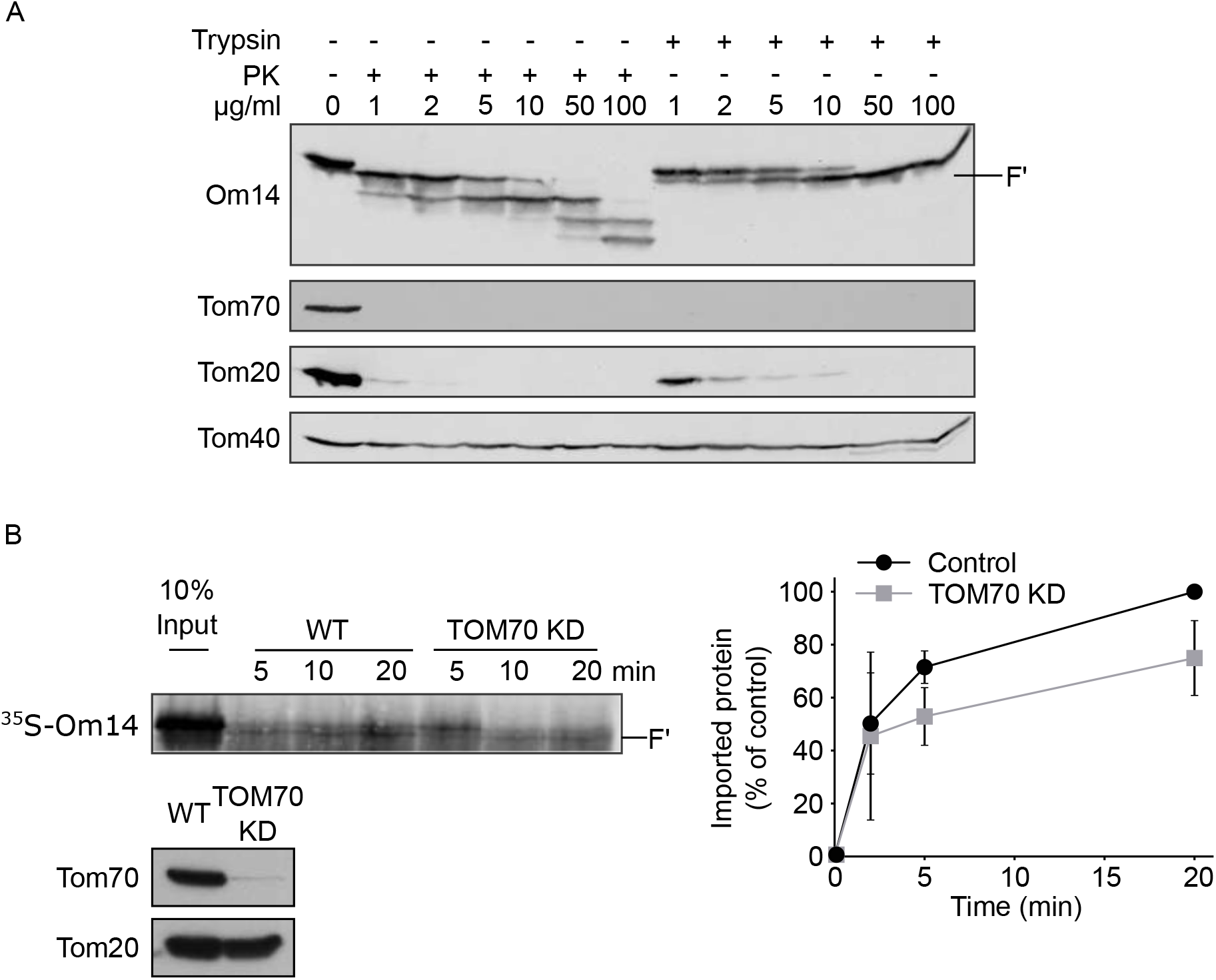
TOM70 contributes to the import of Om14 into mammalian mitochondria. (A) Isolated yeast mitochondria were treated with the indicated concentrations of either Proteinase K (PK) or trypsin. Mitochondrial proteins were then analyzed by SDS-PAGE and immunodecoration with the indicated antibodies. Tom20 and Tom70 are exposed on the surface of the organelle while Tom40 is embedded within the membrane. (B) Radiolabeled Om14 was imported into mitochondria isolated from either control human HeLa cells or HeLa cells where Tom70 was knocked down. After the import, samples were treated with trypsin and analyzed and quantified as described in the legend to Fig. 2B. The lower panel shows analysis of the isolated mitochondria by SDS-PAGE and Western blotting with the indicated antibodies. Note, TOM70 is hardly detected in the KD cells. Right panel: Quantification of the import efficiency was done as described in the legend to Fig. 2B.

**Figure S3.**
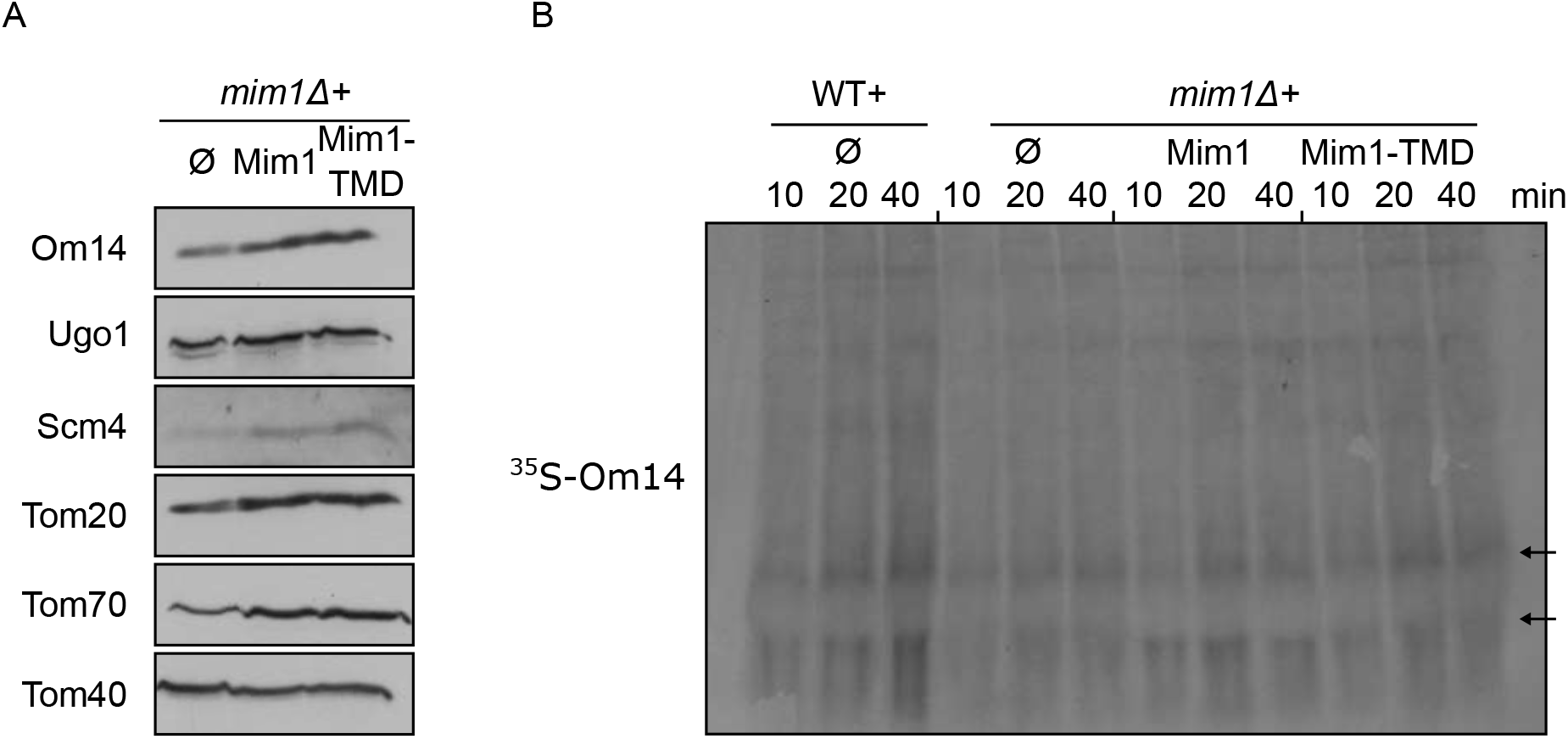
The central TMD of Mim1 can replace the native protein. (A) Mitochondria were isolated from *mim1Δ* cells harboring an empty vector (Ø) or a plasmid encoding either native Mim1 or its central TMD. The steady-state levels of known Mim1 substrates as well as of Tom40 (for comparison) were analyzed by SDS-PAGE and immunodecorated with the indicated antibodies. (B) Radiolabeled Om14 was imported into mitochondria isolated from the indicated cells and further analyzed by BN-PAGE and autoradiography. Both arrows mark oligomeric species of Om14.

**Figure S4.**
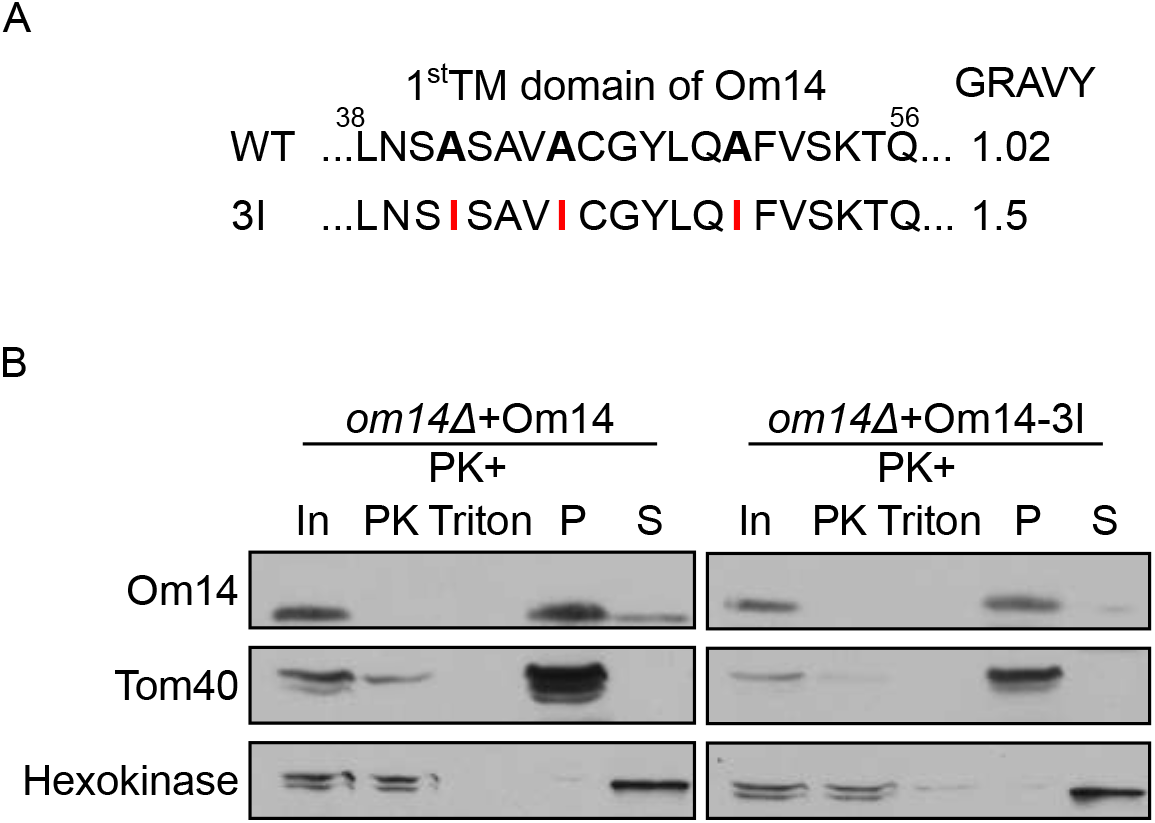
Increasing the hydrophobicity of TMD1 does not affect the membrane integration of Om14. (A) Amino acid sequence of the putative first TMD of both native Om14 (WT) and a variant with increased hydrophobicity due to the introduction of three Ile residues (3I). Mutated residues are in bold. (B) Mitochondria isolated from *om14Δ* cells expressing either native Om14 or the Om14-3I variant were subjected to Proteinase K treatment or alkaline extraction. Samples were further analyzed by SDS-PAGE and immunodecoration with the indicated antibodies.

